## Supplementary figures and images for "Hot springs viruses at Yellowstone National Park have ancient origins and are adapted to their thermophilic hosts"

### Extended_Data_Fig_1

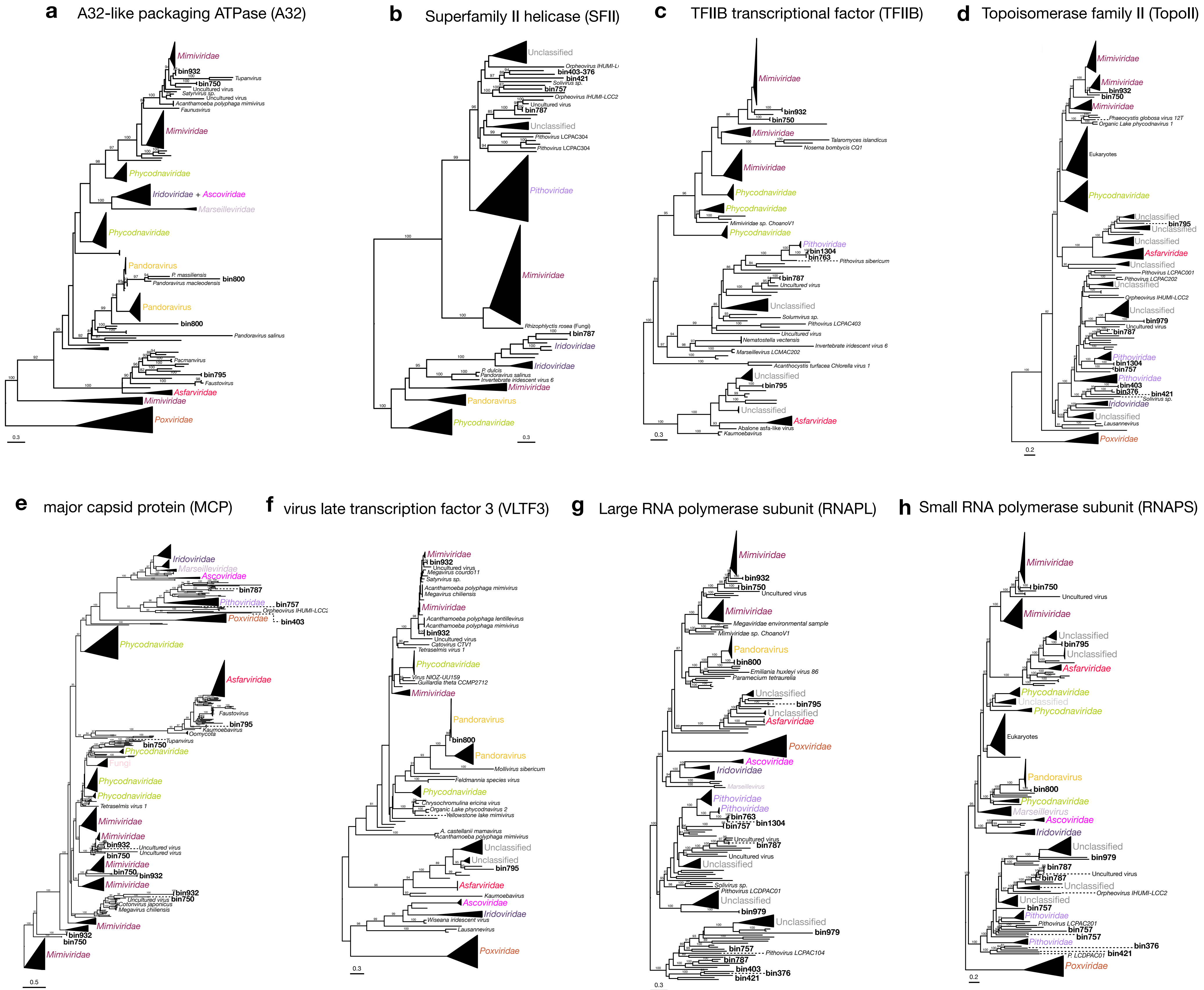
